# Atypical Neurocognitive Visual and Bodily Self-Face Processing in Children with ASD Links to Social Symptomatology

**DOI:** 10.64898/2026.11.19.767776

**Authors:** Tingyu Zhang, Haoyue Qian, Yijia Tang, Siyao Cao, Xiaoyan Zhang, Chenhuan Ma, Shilu Kang, Jieqiong Liu, Neng Gong, Guanghai Wang

**Author notes:** Correspondence to (GHW), (NG) and. These authors contributed equally to this work (Tingyu Zhang, Haoyue Qian, Yijia Tang). **Lead Contact:** Guanghai Wang, PhD, Department of Developmental and Behavioral Pediatrics, Pediatric Translational Medicine Institute, Shanghai Children’s Medical Center, Shanghai Jiao Tong University School of Medicine, Shanghai, China.

## Abstract

**Objective:** To characterize dissociable neural responses in the inferior frontal gyrus (IFG) during visual self-face processing and the temporoparietal junction (TPJ) during bodily self-face processing, and examine their associations with social deficits in children with autism spectrum disorder (ASD).

**Methods:** Real-time self-face videos with and without a red dot were used to establish pure visual-only and integrated visual-bodily self-face processing conditions, respectively. Functional near-infrared spectroscopy (fNIRS) data were acquired from three child groups: typically developing (TD) children, children with ASD who passed self-face recognition (ASDP), and children with ASD who failed self-face recognition (ASDF).

**Results:** In TD children, self-face stimuli activated the left IFG across both conditions, while right TPJ activation was specific to visual-bodily self-face processing. The ASDP group showed preserved left IFG activation but absent right TPJ activation, whereas the ASDF group lacked self-face-specific left IFG activation. Control analyses ruled out attention, exogenous sensory processing, and developmental trajectories as confounds for the diminished right TPJ activation in ASD. Elastic net regression further identified atypical self-face-related neural responses as the primary predictor of reduced social communication proficiency in ASD.

**Conclusion:** Unlike TD children, who engage both neural pathways, children with ASD rely preferentially on left IFG-mediated visual self-recognition while showing impaired TPJ-dependent bodily self-face processing. These atypical neural patterns predict social symptom severity. Our findings reveal novel neurocognitive mechanisms of self-processing and may inform precise assessment and individualized interventions for social deficits in ASD.

**Plain language summary:** Children with autism spectrum disorder (ASD) often struggle with self-face recognition. Using live video and wearable imaging, we examined visual and bodily contributions. Typically developing children activated both systems, whereas ASD children who recognized themselves showed only visual responses and those who did not showed no activation. These findings link altered visual-bodily integration to social deficits, suggesting a mechanism underlying social symptoms and personalized support.

## Introduction

Autism spectrum disorder (ASD) is a complex neurodevelopmental disorder characterized by heterogeneous and multifaceted symptomatology, with social difficulties representing a core area of impairment and a primary focus of intervention (American Psychiatric Association, 2022). A growing body of research has identified self-processing, particularly self-face processing, as a potentially key neurocognitive domain underlying autistic symptomatology (Kaganoi et al. 2025; Kana et al. 2017; Lombardo et al. 2010; Uddin 2011). This capacity underpins fundamental self-other differentiation, laying an indispensable cognitive foundation for children’s social development (Amsterdam 1972; Keromnes et al. 2019; Lombardo and Baron-Cohen 2011). Therefore, elucidating the neurocognitive mechanisms underlying self-face processing in children with ASD is critical for understanding their social deficits and potentially optimizing assessment and intervention strategies.

Accumulating multimodal neuroimaging evidence has revealed atypical neural responses to self-face stimuli in children with ASD relative to typically developing (TD) children. Functional magnetic resonance imaging (fMRI) studies have shown that TD individuals exhibit selective BOLD responses in the middle cingulate cortex during passive self-face viewing, whereas this response is absent in ASD (Lu et al. 2014). Using a more child-friendly technique, functional near-infrared spectroscopy (fNIRS), researchers have reported attenuated activation in the right inferior frontal gyrus (IFG) during self-face recognition in youth with ASD (Kita et al. 2011). Likewise, event-related potential (ERP) studies have identified diminished neural responsiveness to self-face cues in children with ASD, suggesting atypical attentional processing of self-relevant facial information (Cygan et al. 2014; Cygan, Nowicka, and Nowicka 2022). Collectively, these findings suggest that children with ASD exhibit attenuated neural and cognitive responses to their own faces. However, the reported neural patterns underlying this atypical processing remain inconsistent, and the precise neurocognitive mechanisms are yet to be fully elucidated.

Two dominant hypotheses have been proposed to explain aberrant self-face processing in ASD, each highlighting distinct domain-specific functional impairments: one attributes the abnormality to dysfunction in visual self-processing cortical regions (e.g., the IFG), while the other emphasizes abnormal neural activity in regions mediating bodily self-face perception (e.g., the TPJ). Decades of research have delineated that typical self-face recognition relies on two functionally dissociable yet complementary neural pathways (Sugiura 2015). The first mediates visual self-recognition based on long-term stored facial visual representations; the second subserves bodily self-face perception via real-time multisensory body-related processing (Apps et al. 2015). The latter mechanism underlies the classic mirror mark-test paradigm (Amsterdam 1972; Gallup 1970), and real-time dot-overlaid self-face stimuli can engage both pathways, enabling the experimental dissociation of visual and visual-bodily integrated self-processing. At the neural level, two meta-analyses have converged on a core self-face processing network, including bilateral IFG, bilateral insula, right postcentral gyrus, right anterior cingulate cortex (ACC), and bilateral TPJ (Hu et al. 2016; Qin, Wang, and Northoff 2020). Specifically, the left IFG and ACC predominantly support visual and conceptual self-recognition, whereas the TPJ and insula are closely linked to bodily self-perception. Accordingly, attenuated self-face neural responses in children with ASD may stem from either dysfunction in visual self-recognition regions (e.g., IFG) or deficits in bodily self-processing regions (e.g., TPJ), which require further investigation. This study therefore aims to directly characterize dissociable neural responses in children with ASD relative to TD.

Beyond neural characterization, the association between self-face processing and typical social development has long been recognized (Amsterdam 1972; Keromnes et al. 2019; Lombardo and Baron-Cohen 2011). It has been proposed that atypical neurocognitive self-processing undermines social competence in individuals with ASD, yet this hypothesis requires further scrutiny, particularly given the dissociation between visual and bodily mechanisms. Recent work has demonstrated that fNIRS is well suited for capturing neural activity in this population due to its tolerance for motion artifacts and high wearability (Liu et al. 2019). Atypical brain activity in ASD also shows potential as a predictive biomarker for ASD functional performance assessment (Kou et al. 2026; Liu et al. 2026), and mapping aberrant neurocognitive processing of self-faces may further assist clinicians in comprehensive ASD evaluations. Accordingly, a secondary aim of this study is to examine whether atypical neurocognitive self-face processing can predict social performance in ASD.

In summary, this study used fNIRS to examine dissociable neural responses in the IFG (visual self-face processing) and TPJ (visual-bodily integrated self-face processing) in children with ASD, and their predictive value for core social deficits. Real-time self-face stimuli with or without a red dot were applied to elicit the two processing modes across three groups: TD children, children with ASD who passed self-face recognition (ASDP), and those who failed (ASDF). We first compared TD and ASDP groups to identify ASD-specific atypical activation, and compared ASDP and ASDF groups to verify the neural specificity of self-recognition in ASD. Subsequently, we confirmed that the neural pattern observed above stemmed from atypical self-face processing rather than variations in attentional allocation, exogenous sensory processing, or neurodevelopmental trajectories. Finally, machine learning was used to test whether self-face-related activation predicts social symptomatology in ASD (see Figure 1).

**Figure 1:**
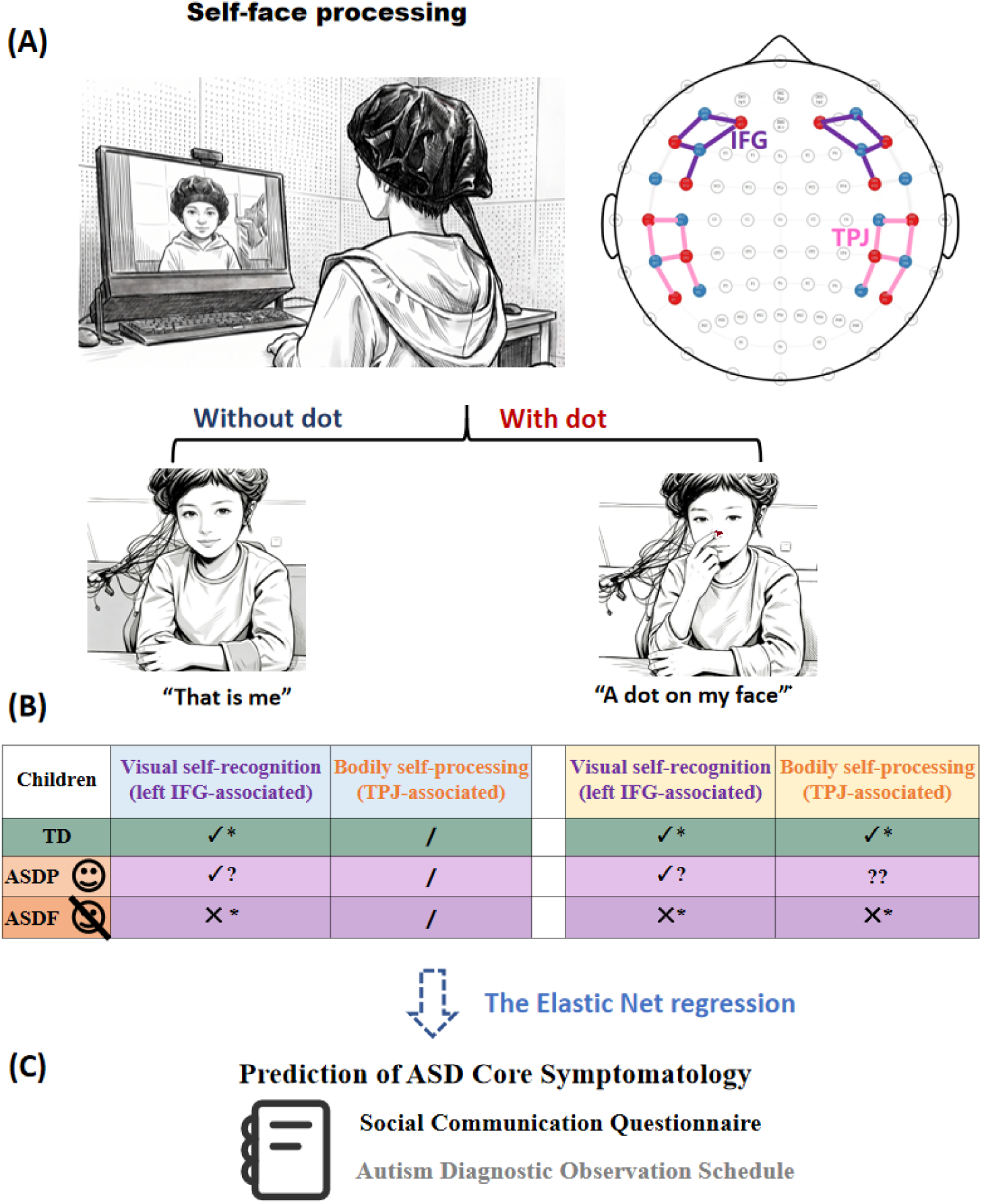
Study design and hypothesis. (A) The specific neurocognitive patterns underlying self-face processing in children with ASD were examined by considering two main dissociable neural responses to self-face separately, one during the face-without-dot condition and the other during the face-with-dot condition (which involves the bodily self). fNIRS recording was used to focus on the activities of the bilateral IFG and TPJ. (B) We hypothesized that TD children would show a selective response of the left IFG to visual self-face, together with bodily self-related right TPJ activation, and that these responses would be absent in ASD children who failed self-face recognition (ASDF). The neural response pattern in ASD children who passed self-face recognition (ASDP) remained to be tested. (C) We further predicted that atypical self-face processing would predict the core symptomatology in ASD, using elastic net regression.

## Methods

### Participants

From September 2023 to September 2025, an initial cohort of 204 children was recruited via online and offline advertisements. Following rigorous clinical screening and data quality verification, the final analytical sample consisted of 50 children with ASD and 30 age-matched TD children, all of whom completed all experimental tasks and behavioral assessments. The ASD sample was divided by self-face recognition performance into 20 children who succeeded (ASDP) and 30 who did not (ASDF).

Participants were aged 2–7 years with normal or corrected-to-normal vision and intact motor function. ASD participants had a confirmed DSM-5 diagnosis by a developmental pediatrician and underwent ADOS-2 assessment by research-reliable clinicians. TD participants had no history of developmental delay, neurological or psychiatric disorders. All participants were excluded for severe systemic/organic illness requiring hospitalization, a history of epilepsy or organic brain injury, a significant familial psychiatric history, or current/prior long-term psychotropic medication use. The detailed recruitment flowchart is shown in Figure S1.

The study protocol was approved by the Institutional Review Board of the Shanghai Children’s Medical Center (Approval No. SCMCIRB-K2022187-1) prior to any data collection. Written informed consent was secured from the parents or legal guardians of all participants before study commencement.

A priori power analysis (assuming a small effect size *f* = 0.15, *α* = 0.05, and 80% power) indicated a minimum requirement of 63 participants. Our final analyzable sample of 80 participants exceeded this threshold (see Methods S1 for details).

### Clinical and Main Symptoms Assessments

#### ASD diagnoses and symptom

Clinical diagnoses were established through a comprehensive evaluation, incorporating semi-structured caregiver interviews based on DSM-5 criteria and direct behavioral observation with ADOS-2 (Gotham et al., 2008). The ADOS-2 was administered using the Toddler module or Modules 1–4 selected according to each child’s chronological age and expressive language ability. To quantify core ASD symptom severity and enable comparisons across ADOS-2 modules, Calibrated Severity Scores (CSS; 1–10) were derived for overall symptoms, Social Affect (SA CSS), and Restricted and Repetitive Behaviors (RRB CSS), indexing dimensional symptom variation.

#### Social functioning assessment

Parents completed the Social Communication Questionnaire (SCQ) to assess children’s social communication abilities based on their behavior over the preceding three months. The 40 dichotomous (Yes/No) items assess social interaction, communication, and restricted and repetitive behaviors. Items indicative of atypical behavior are scored as 1 and summed to generate a total score (range: 0–40), with higher scores reflecting greater social communication difficulties (Rutter 1999).

### Task and procedure

#### Face processing paradigm

According to our research design, face stimuli were presented under two experimental conditions: face-only presentation and face-with-red-dot presentation, yielding four experimental phases: self-face only, unfamiliar-face only, self-face with a red dot, and unfamiliar-face with a red dot (see Figure S2A). A 10-second fixation cross was displayed prior to each phase to serve as the baseline.

In the face-only condition, participants viewed a 1-minute live video of their own face (the self-face visual processing task) and a 1-minute pre-recorded video of an age-and sex-matched unfamiliar child’s face. The two types of face stimuli were presented in pseudorandom order across all participants. Immediately following each video presentation, the experimenter pointed to the screen and asked the participant, “Who is this?” to elicit a verbal identification response. In addition, verbal cues and gestures were used when necessary to sustain participants’ attention throughout the task.

In the face-with-red-dot condition, participants watched a 1-minute live video of their own face overlaid with a red dot, mimicking the classic mirror mark-test paradigm. This setup engages both visual and bodily self-processing, and may trigger spontaneous tactile exploration toward the dot, which defines this phase as the visual-bodily self-face processing condition. For the control contrast, participants also viewed a 1-minute pre-recorded unfamiliar-face video with an identical red dot overlay. The pseudorandom stimulus sequence and attention-maintaining protocols adopted in this condition were identical to those used in the face-only condition.

#### Experimental procedure

During the formal experiment, participants sat approximately 60 cm from a computer monitor. A Logitech C920Pro 1080p webcam delivered real-time visual feedback, horizontally flipped via PotPlayer to mimic a mirror reflection.

Prior to the face-only task, each child watched a 1-minute resting-state animation to calm down. Another 1-minute resting animation was played during which the experimenter secretly placed an odorless, non-irritating red dot of face paint on the tip of the child’s nose. The experiment was terminated if the child noticed this marking procedure, after which the face-with-red-dot task was administered (see Figure S2B).

We employed a dual-computer setup: one computer presented stimuli via Tobii Pro Lab and transmitted synchronization triggers, while the second recorded fNIRS signals.

#### Self-face recognition behavioral assessment

Upon completion of the formal experimental paradigm, participants completed a validated tissue test designed to verify children’s self-recognition performance. All verbal and nonverbal experimental prompts were withdrawn in this phase: the experimenter merely handed a tissue to each child and observed spontaneous wiping behaviors directed toward the facial mark. No fNIRS data were collected during this test.

Self-recognition success was operationalized as the emergence of self-directed behaviors captured either during the red-dot self-face recording session or the tissue test, including tactile contact with or deliberate wiping of the facial mark and verbal self-identification.

### fNIRS data acquisition and analysis

Cortical hemodynamic signals during face processing were recorded using a portable continuous-wave fNIRS system (NIRSport2, NIRx Medical Technologies, USA). Measurements focused on a priori defined regions of interest (ROIs): the bilateral IFG and TPJ. Preprocessing and statistical analyses were conducted in MATLAB 2022b using the HOMER2 (Huppert et al. 2009) and NIRS-SPM packages (Ye et al. 2009). Detailed procedures regarding optode placement (Yucel et al. 2021), data preprocessing workflows (Fishburn et al. 2019; Whiteman et al. 2018) and GLM specifications (Worsley and Friston 1995) are provided in the Supplementary Material (Methods S2, Table S1, and Figure S2).

### Statistical analyses

#### Linear Mixed-Effects Modeling for brain activation analysis

To characterize group-specific task-evoked brain activation patterns, linear mixed-effects models (LMMs) were separately applied to the three participant groups: TD, ASDP, and ASDF.

Prior to model fitting, an outlier detection procedure was applied to the ROI activation data (left/right IFG, left/right TPJ). For each participant and each experimental condition, activation values exceeding 3 standard deviations from the conditional mean were excluded from subsequent analysis to eliminate extreme value interference.

LMMs were adopted to analyze condition-related β-values, accounting for the nested data structure and inter-individual variability. Separate LMMs were built for each ROI. The models included face type (self vs. unfamiliar other) and red dot presentation (present vs. absent) as categorical fixed effects, chronological age as a continuous covariate, and subject-specific random intercepts to control for repeated measures. The model formula was: Activation ∼ Face Type × Red Dot + Age + (1 | Subject).

All LMMs were fitted via restricted maximum likelihood (REML) estimation with a full Cholesky covariance structure and effect coding for categorical variables. Standardized regression coefficients, t-statistics, degrees of freedom, and p-values were calculated via the Satterthwaite approximation to evaluate fixed-effect significance. To explore the fine-grained spatial basis of regional activation effects, the same LMM framework was applied to individual channel-level data. The Benjamini-Hochberg false discovery rate (FDR) correction was performed on channel-level p-values of the main self-face effect within each ROI to strictly control multiple comparison bias while preserving statistical power.

#### Self-Face Processing Index Analysis

A self-face processing index was defined for each ROI as the hemodynamic activation contrast between self-face and other-face conditions. Group differences in self-face processing indices were examined via independent GLMs, with group (i.e., TD/ASDP/ASDF) as the fixed factor and grand-mean-centered chronological age as the continuous covariate to eliminate age-related baseline differences and multicollinearity. The model formula was: Activation ∼ Group + Age.

Omnibus F-tests were first performed to detect global group main effects. Pre-specified planned pairwise comparisons were then conducted based on group activation characteristics. Age-adjusted estimated marginal means (EMMs) were calculated, and one-sample t-tests against zero were performed for each group. All comparative results were reported with parameter estimates, p-values, and Cohen’s d effect sizes.

#### Confounding Factor Control Analysis

To exclude the confounding effects of attentional bias induced by red dot stimuli, an LMM was used to test whether red dot presentation modulated right TPJ activation during other-face processing across the three groups, with the formula: Activation ∼ Group × Red Dot + Age + (1 | Subject).

To further rule out tactile sensory interference, we focused on the ASDF group and compared right TPJ activation between children with and without spontaneous tactile touching behaviors during red dot presentation, under both self-face and other-face red dot conditions.

#### Atypical Developmental Trajectory Analysis

For the developmental trajectory analysis, all participants were divided into TD and ASD groups. Missing clinical and activation data were imputed via the median substitution method, and all brain activation features were standardized prior to modeling.

Quadratic polynomial regression was performed on standardized activation data to identify atypical neurodevelopmental patterns in the ASD group. The regression model included fixed effects of group (TD vs. ASD), linear age, quadratic age (Age²), and group-by-age linear/quadratic interaction terms to capture nonlinear developmental trends: Activation ∼ Group × Age + Group × Age².

### Machine learning analyses approach

We used Elastic Net regression to predict behavioral phenotypes from baseline-adjusted self-other activation contrasts within predefined ROIs. SCQ scores were log-transformed to reduce skewness, whereas ADOS comparison scores were analyzed on their original scale. The mixing parameter was fixed at α = 0.1, combining *L1* (Lasso) variable selection and *L2* (Ridge) regularization.

A 20-repeated five-fold cross-validation strategy was applied to improve model generalizability and reduce sampling variability. To prevent information leakage, all preprocessing steps were performed independently within each training fold. Missing neuroimaging features and age values were imputed using training-set medians. Neuroimaging features were residualized for chronological age and standardized using training-derived parameters, which were then applied to the corresponding held-out fold. Within each training fold, the regularization parameter (λ) was optimized using three-fold cross-validation by minimizing mean squared error. The optimized model was subsequently applied to the held-out fold to generate out-of-sample predictions, which were averaged across repetitions to obtain participant-level predicted scores.

Model predictive performance was then evaluated using the R² indicator: Pearson’s or Spearman’s correlation coefficients between observed and predicted scores, cross-validated predictive coefficient of determination (predictive R²).

Feature importance was defined as the mean absolute regression weight across all cross-validation folds, and model stability was assessed via feature selection frequency. Features were ranked by importance to identify the core neural biomarkers corresponding to behavioral phenotypes. To evaluate whether model prediction performance significantly exceeded chance levels, 1,000 non-parametric permutation tests were performed to derive empirical *p*-values (Phipson and Smyth 2010).

Detailed machine learning procedures are provided in Supplementary Methods S3.

## Results

### Demographic information and clinical characteristics

As listed in Table 1, there were no significant group differences in gender or age across the three groups. ADOS general symptomatology, based on ADOS-2, did not differ significantly between the ASDP and ASDF subgroups. Of note, developmental age (DA) differed significantly between the TD and ASD groups; accordingly, age was treated as an important confounder across all analyses in this study.

**Table 1.** Participant demographic information and clinical characteristics.

|  | TD (N=30) | ASD (N=50) | ASDP (N=20) | ASDF (N=30) | TD vs ASD | ASDP vs ASDF |
|---|---|---|---|---|---|---|
| **sex, N males** | 22 (73.3%) | 45 (90.0%) | 19 (95.0%) | 26 (86.7%) |  |  |
| **Age, years** | 5.11 ± 1.32 | 5.57 ± 1.41 | 6.06 ± 1.17 | 5.24 ± 1.48 |  |  |
| **SA CSS** | - | 6.50 ± 1.63 (n=50) | 6.30 ± 1.45 (n=20) | 6.63 ± 1.75 (n=30) |  |  |
| **RRB CSS** | - | 5.72 ± 1.97 (n=50) | 5.15 ± 2.21 (n=20) | 6.10 ± 1.73 (n=30) |  |  |
| **CSS** | - | 6.12 ± 1.53 (n=50) | 5.90 ± 1.62 (n=20) | 6.27 ± 1.48 (n=30) |  |  |
| **Developmental Age (DA), months** |  |  |  |  |  |  |
| Locomotor DA | 70.11 ± 18.11 (n=27) | 50.51 ± 14.28 (n=40) | 53.00 ± 10.31 (n=13) | 49.31 ± 15.88 (n=27) | *** |  |
| Personal-Social DA | 67.35 ± 15.81 (n=27) | 38.16 ± 10.62 (n=40) | 43.42 ± 11.39 (n=13) | 35.63 ± 9.42 (n=27) | *** | * |
| Hearing and Language DA | 73.77 ± 17.86 (n=26) | 34.70 ± 12.64 (n=40) | 40.04 ± 7.60 (n=13) | 32.13 ± 13.85 (n=27) | *** |  |
| Eye-Hand Coordination DA | 64.63 ± 18.44 (n=27) | 45.90 ± 15.99 (n=40) | 53.81 ± 16.52 (n=13) | 42.09 ± 14.54 (n=27) | *** | * |
| Performance DA | 68.94 ± 17.47 (n=27) | 46.96 ± 17.04 (n=40) | 57.27 ± 18.99 (n=13) | 42.00 ± 13.80 (n=27) | *** | ** |
| Practical Reasoning DA | 75.86 ± 13.91 (n=21) | 46.88 ± 20.67 (n=23) | 53.00 ± 19.41 (n=9) | 42.95 ± 21.18 (n=14) | *** |  |

*Note.* ADOS-2 = Autism Diagnostic Observation Schedule, Second Edition; CSS = Calibrated Severity Score; SA = Social Affect; RRB = Restricted and Repetitive Behaviors. Values represent Mean ± SD unless specified otherwise. Group differences were assessed using two-sided independent-samples t-tests for continuous variables and chi-square tests for categorical variables. Asterisks indicate uncorrected p values: \*p < 0.05, \*\*p < 0.01, and \*\*\*p < 0.001.

### The IFG and TPJ responses to visual-only and visual-bodily self-face processing conditions across three children groups

We first fitted LMMs to investigate face-processing-related hemodynamic activation in the bilateral IFG and TPJ across the three groups. Face type (self vs. other) and red dot (with vs. without) were defined as fixed predictors, while chronological age was included as a confounding covariate to account for inter-individual age variability. In the TD group, significant self-face processing effects were observed in the left IFG and right TPJ. Specifically, the right TPJ showed a significant main effect of face type (p = 0.021, Cohen’s d =-0.266) and a significant face type × red dot interaction effect (p = 0.016, Cohen’s d = 0.279), with stronger activation for self-face (vs. other-face) under the red dot condition. The left IFG exhibited a marginally significant main effect of self-face processing (p = 0.078, Cohen’s d =-0.196). Further channel-level analysis with intra-ROI FDR correction confirmed this reliable trend, revealing significantly enhanced activation in channel 2 (FDR-corrected p = 0.035) and channel 4 (FDR-corrected p = 0.048) in response to self-face stimuli (Figure 2A).

**Figure 2:**
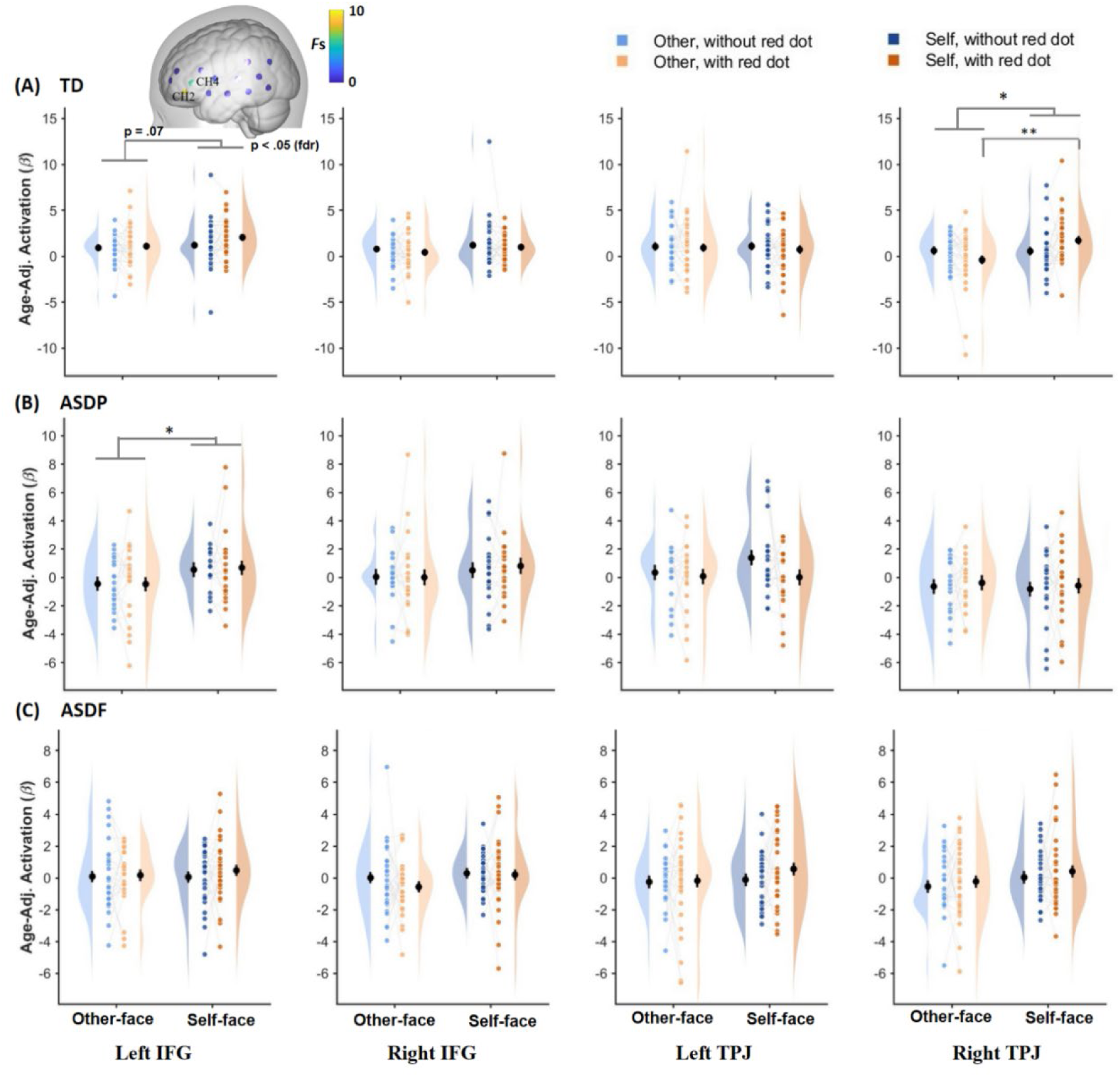
Activations in the IFG and TPJ during self-face versus other-face processing across three groups of children. In TD children, a selective response of the left IFG to self-face (vs. other child’s face) was observed in both the without-dot (visual only) and with-dot (visual-bodily) conditions, whereas enhanced right TPJ activation to self-face was observed only in the with-dot condition. In the ASDP group, only the selective left IFG response to self-face was found in both conditions. In the ASDF group, no selective response to self-face was observed in the bilateral IFG and TPJ. Note: * p < 0.05, ** p < 0.01

Differing from the TD group, the ASDP group exhibited a significant main effect of face type in the left IFG (p = 0.026, Cohen’s d =-0.308), with heightened activation for self-face (vs. other-face). No significant main effects of face type or red dot presentation, nor significant interaction effects, were observed in the right IFG or bilateral TPJ in the ASDP group (all ps > 0.05; Figure 2B).

By contrast, the ASDF group exhibited no significant main effects of self-face processing, red dot presentation, or age, and no significant two-way interactions in all four predefined ROIs (all ps > 0.05; Figure 2C).

Group differences in self-face processing patterns were further characterized using the self-face processing index. Significant left IFG activation during general visual self-face processing (across with red-dot and without red-dot conditions) was identified in the TD group, t(73) = 2.655, p = 0.010, Cohen’s d = 0.501, and the ASDP group, t(73) = 2.339, p = 0.022, Cohen’s d = 0.414, but not in the ASDF group, t(73) = 1.252, p = 0.215, Cohen’s d = 0.334; see Figure 3A. Meanwhile, significant right TPJ activation specific to visual-bodily self-face processing (with red-dot condition only) was exclusively observed in TD children, t(73) = 2.279, p = 0.026, Cohen’s d = 0.420, whereas neither ASD subgroup showed comparable activation (all ps > 0.05; Figure 3B).

**Figure 3:**
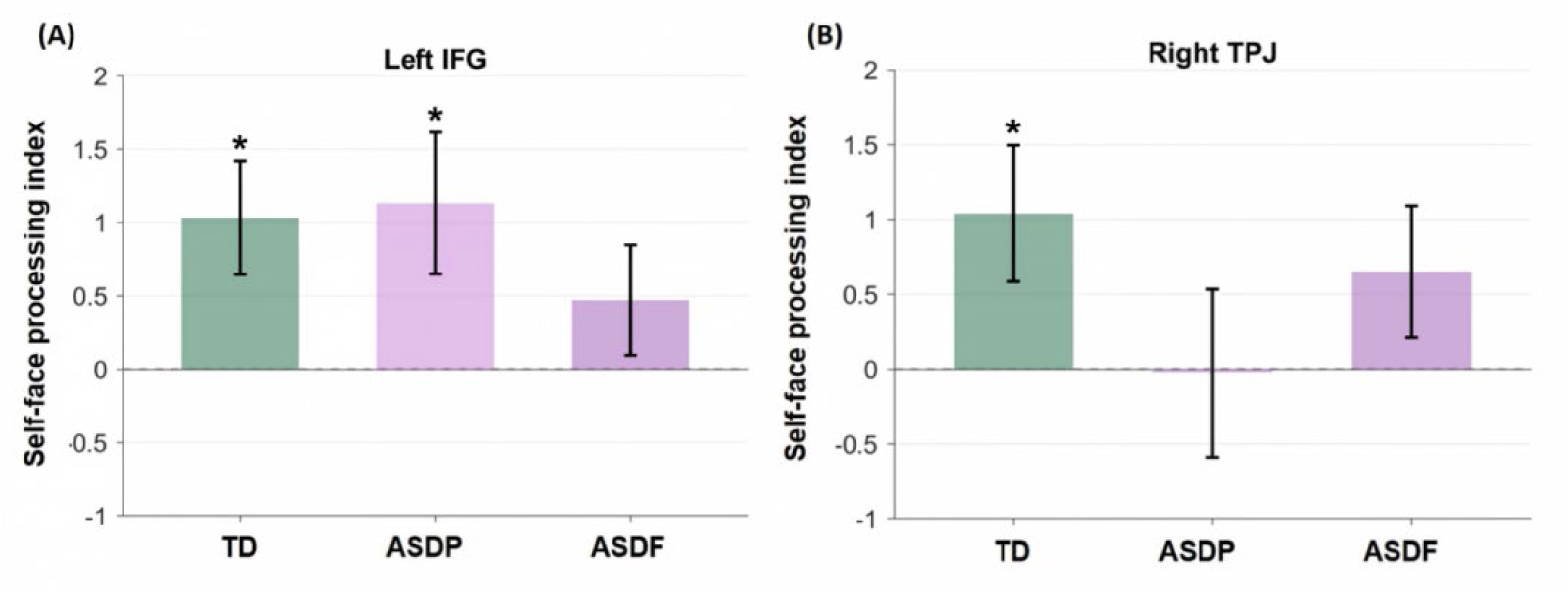
Group differences in the self-face processing index. A significant self-face processing effect (βself – βother) of the left IFG across both experimental conditions (face with and without the red dot) was observed in the TD and ASDP groups, but not in the ASDF group. A significant self-face processing index of the right TPJ for the visual-bodily condition (face with red dot) was detected only in the TD group (p < 0.05).

Analyses of the remaining predefined ROIs revealed no significant activation differences across groups (all p > 0.05). Full statistical results from all analyses are available in the Supplementary Materials (Table S2–S4).

### The atypical neural pattern of self-face processing in ASD is independent of attentional allocation, external sensory processing, and developmental delay

The above results indicated that children with ASD retained left IFG activation during self-face recognition but showed absent right TPJ activation during visual-bodily self-face processing. To verify whether the diminished TPJ response stemmed from atypical attention allocation, impaired external sensory processing, or generalized developmental delay in ASD, we conducted a series of confounding factor exclusion analyses.

First, we tested whether red dot stimuli altered domain-general attention and further modulated right TPJ activation during face processing. To evaluate this, an LMM analysis was conducted in the other-face condition, which revealed no significant main effect of red dot presentation, F(1, 68.969) = 0.189, *p* = 0.665, η_p_^2^ = 0.003, and no significant group × red dot interaction, F(1, 68.975) = 1.888, p = 0.159, η_p_*^2^ = 0*.052 (Figure 4A).

**Figure 4:**
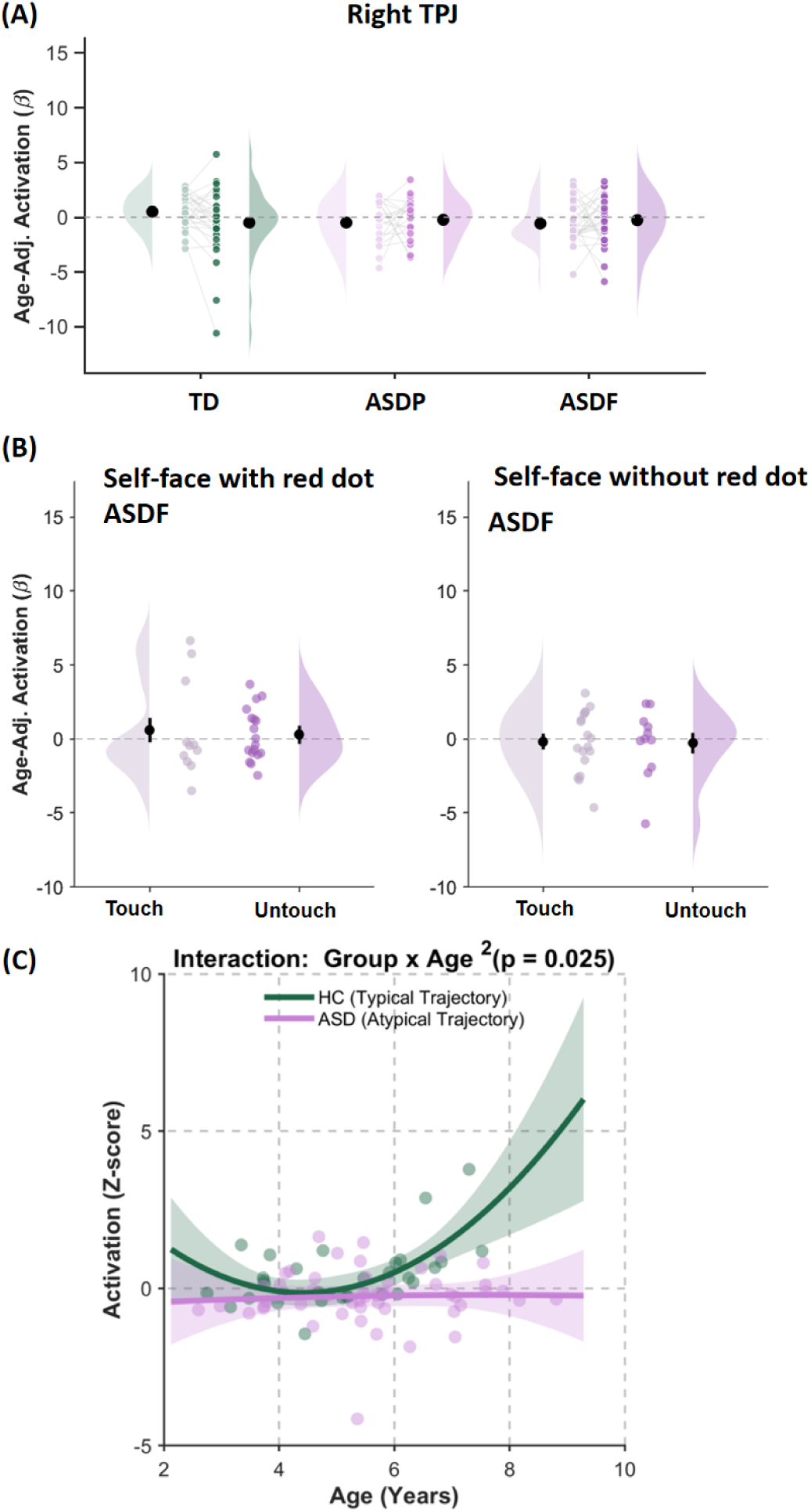
Control analysis of the atypical neural pattern of self-face processing in ASD. The atypical neural pattern of self-face processing observed in individuals with ASD was independent of three confounding variables: (A) attentional allocation, as evidenced by the absence of a dot presentation effect in the other-face condition; (B) external sensory processing, as indexed by no significant group difference between the touch and no-touch subgroups within the ASDF cohort; (C) developmental delay, as indicated by a non-significant age-related effect in the ASD group.

Second, we ruled out potential confounding effects of exogenous tactile sensory input elicited by red-dot touching behaviors. Within the ASDF group, approximately half of the children exhibited spontaneous facial touching during red-dot presentation, whereas the remaining participants did not. We therefore compared right TPJ activation between ASDF children with and without spontaneous nasal dot-touching behavior under both self-face and other-face red-dot conditions. No significant group differences in activation were observed in either condition, self-face: F(1, 26) = 0.076, p = 0.784, η_p_^2^ *=* 0.003; other-face: F(1, 26) = 0.008, p = 0.931, η_p_^2^ < 0.001; see Figure 4B.

Third, quadratic polynomial regression was used to explore whether atypical neurodevelopmental trajectories accounted for the absent right TPJ response in ASD. The results revealed diagnosis-dependent nonlinear developmental trajectories specifically in the right TPJ during self-face processing with a red dot. The linear age × group interaction showed a marginal trend toward significance (p = 0.055), and the quadratic age² × group interaction reached statistical significance (p = 0.025). Developmental curve analysis showed that TD children exhibited a typical nonlinear developmental pattern: right TPJ activation gradually decreased before 4 years of age and subsequently increased. In contrast, children with ASD showed flattened and attenuated right TPJ developmental trajectories (Figure 4C).

Collectively, these exclusion analyses confirm that the attenuated right TPJ response during visual-bodily self-face processing in children with ASD reflects inherent atypical self-representation neural mechanisms, rather than group differences in attentional allocation, external tactile sensory processing, or generalized neurodevelopmental delay.

### The atypical neural processing of Self-face and Its Prediction of Core Social Symptoms in ASD

Using the primary Elastic Net regression framework (with the mixing parameter fixed at α = 0.10), we examined whether atypical neural processing patterns underlying self-face representation in ASD significantly predicted children’s core phenotypic traits, particularly social impairments. The predictive models yielded significant performance for both general social communication deficits and core autism symptomatology. For the SCQ total score (group-level effects are presented in Table S5), the model produced a robust Pearson correlation between observed (log-transformed) and predicted scores of r = 0.381 (prediction p < 0.001; permutation p = 0.009, Figure 5A). The model also demonstrated significant predictive accuracy, with a mean absolute error (MAE) of 0.548 (permutation p = 0.010) and a cross-validated predictive R² of 0.145 (permutation p = 0.009; Figure 5B).

**Figure 5:**
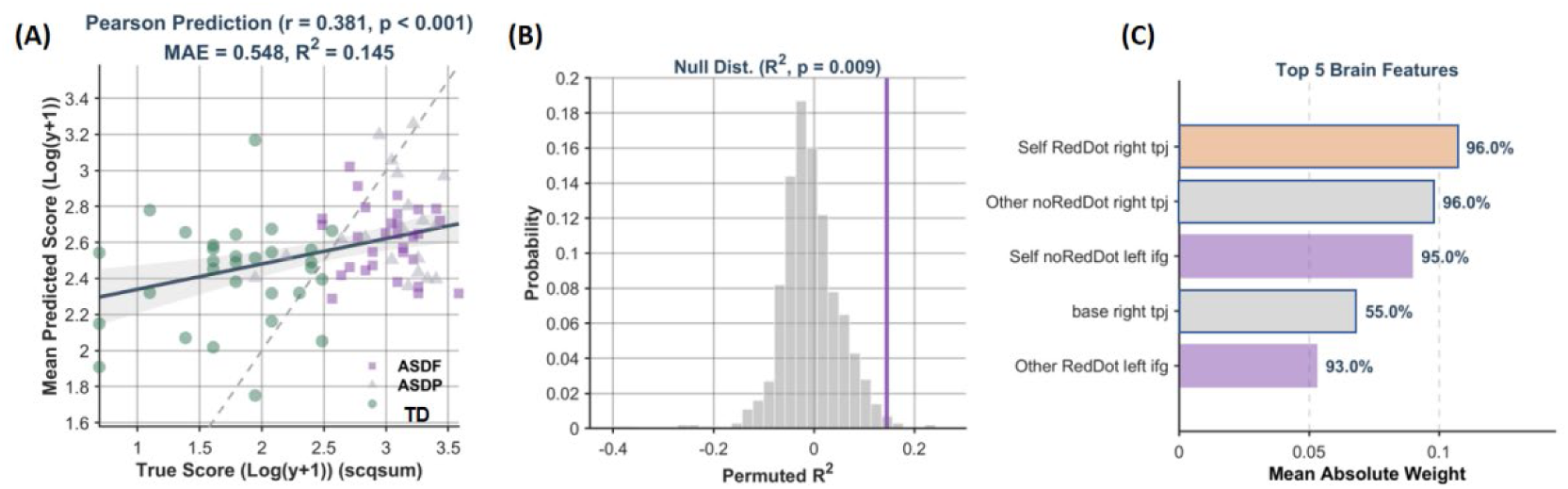
Elastic Net Regression Analysis of Brain Activation Predicting Social Symptoms in ASD. Neural activity during face processing significantly predicted core social symptoms in individuals with ASD (A). This predictive effect survived permutation testing (B). The top five contributing brain features were further extracted and visualized in a panel that included the bodily-self-related right TPJ and the visual-self-related left IFG (C).

Consistent with our hypotheses, neural signatures associated with atypical self-face processing in ASD, including bodily self-related activity in the right TPJ and visual self-face-related activity in the left IFG, emerged as the dominant and most stable predictors (Figure 5C). Intriguingly, TPJ neural activity related to face processing that encompasses both self and other conditions was the predominant predictor of the RRB CSS (prediction p < 0.001; see Figure S3), further supporting the critical role of self-face neural processing in characterizing ASD phenotypes.

## Discussion

Self-face processing is critical for self–other differentiation and social development, especially in children with ASD, yet its neural basis and link to social functioning remain unclear. We experimentally dissociated IFG (visual) and TPJ (bodily) responses to self-face in ASD versus TD children. TD children showed sustained left IFG activation across conditions and right TPJ activation selectively during visual-bodily self-face processing. ASD children who passed self-face recognition (ASDP) retained left IFG activation but lacked right TPJ activation in the visual-bodily condition; those who failed (ASDF) showed no left IFG activation to self-face. Control analyses ruled out attentional, sensory, or developmental confounds. Elastic Net regression revealed that self-face-specific activation predicted social symptom severity in ASD. These findings illuminate neurocognitive mechanisms underlying atypical self-processing and social deficits in ASD.

### Atypical neural responses toward self-face in children with ASD

Our results revealed that during self-face processing, TD children exhibited activation in the left IFG associated with visual self-recognition and the right TPJ linked to bodily self-perception. This observation aligns with prior neuroimaging investigations of self-face processing among neurotypical individuals (Hu et al. 2016; Sugiura 2015). However, this activation of right TPJ associated with self-face has been absent in ASD. Additional control analyses verified that the absence of right TPJ activation was specific to atypical self-face processing, rather than driven by confounding variables such as altered attentional allocation, exogenous sensory processing, or developmental delays in autistic children.

Impaired bodily self-face processing in ASD has also been documented in previous body illusion studies targeting this clinical population. Existing research indicates that autistic individuals exhibit difficulties integrating body-related multisensory cues, which constrains perceptually grounded self-processing (Greenfield et al. 2015; Ropar et al. 2018). Notably, the work of Deltort and colleagues demonstrated that deficient multisensory integration reduces perceptual sensitivity to the physical features of one’s own face in children with ASD (Deltort et al. 2022).

On the other hand, a selective response of the left IFG to self-face was observed in individuals with ASD who passed self-face recognition at a level comparable to TD children, but was absent in those who failed self-face recognition. Considering that the left IFG has been established as a core region for processing self-identity information, including self-face and one’s own name (Qin et al. 2020), our results suggest that some key functional brain regions supporting high-level self-representation (such as the IFG) are preserved in a subgroup of individuals with ASD, thereby supporting their general self-related processing capabilities.

In short, we found that the neural response selectively associated with the bodily self is absent in ASD, whereas neural responses to the high-level self (e.g., identity or concept) are preserved in some individuals with ASD. This observation is in line with previous studies (Kaganoi et al. 2025; Ropar et al. 2018) and provides further evidence from neural activity.

### The role of atypical self-face processing in social functioning in ASD

Beyond characterizing the atypical neural processing pattern underlying self-face processing in ASD, another aim of this study was to directly examine its influence on daily social functioning. While heavily theorized (Suddendorf and Whiten 2001; Tordjman et al. 2019), empirical tests of this association are rare. We found that children’s social functioning deficits, particularly those related to ASD severity, could be primarily predicted by atypical neural responses to self-face stimuli, including the absent activation of the right TPJ as well as preserved IFG functional activity. These findings provide experimental evidence for the association between atypical self-processing and deficits in social function in ASD. The association described above also holds clinical importance for the assessment and intervention of ASD. Specifically, deficits in social function represent one of the core symptoms of ASD. However, previous ASD treatment and assessment mainly (or even merely) focused on cognitive and neural responses to social agents. For example, one of the most widely adopted clinical assessments for ASD, the ADOS-2 (Gotham et al. 2008), does not include any assessment of self-processing. Our findings suggest an important role of self-face processing in the social function deficits observed in ASD, and indicate that ASD assessments should incorporate a self-processing component, especially for young children.

### Limitations and Conclusions

Several limitations should be acknowledged. First, females were underrepresented in our sample, although this imbalance reflects the established male predominance in pediatric ASD. Future studies are therefore required to replicate the current findings in female cohorts and verify the generalizability of these findings for girls with ASD. Second, the current experimental design did not include close-other face stimuli, a stimulus category repeatedly demonstrated to modulate self-processing performance in autistic children. Subsequent research incorporating familiar-face trials would help to systematically characterize the neural discrepancies between self-face and familiar-other face processing in this population.

Despite these limitations, our findings indicate that children with ASD, unlike TD children, engage only the left IFG related to visual self-processing during self-face processing. This atypical neural pattern significantly predicts their social functioning deficits, offering neurocognitive insights that may inform targeted interventions.

## Acknowledgments

This research was funded by the National Science and Technology Innovation 2030 “Brain Science and Brain-inspired Research” Major Project of China (2021ZD0204200), National Natural Science Foundation of China (82401803, 82571607), Shanghai Municipal Health Commission (2022XD056), Shanghai Key Laboratory of Child Brain and Development (24dz2260100), and the Phase II Construction of Shanghai High-Level Local University Innovation Team from Shanghai Jiao Tong University School of Medicine.

## Supplemental materials

### Methods S1

#### Sample size determination

Sample size requirements were determined a priori based on the primary hypothesis that hemodynamic responses would differ across conditions. Because exact prospective power calculations for linear mixed-effects models (LMMs) incorporating repeated observations, continuous covariates, and random effects are computationally challenging, a conservative approximation was conducted using G*Power 3.1 based on a repeated-measures framework.

The power analysis was conducted assuming a small effect size (f = 0.15) for a within-subject condition effect, corresponding to the repeated-measures structure of the experimental design (2 face types × 2 red dot conditions). With an alpha level of 0.05, statistical power of 0.80, and a moderate correlation among repeated measurements (r = 0.50), the estimated minimum total sample size was 63 participants. The final analyzable sample consisted of 80 participants (TD, ASDP, and ASDF groups), exceeding the a priori estimate. This sample size was considered adequate for testing the primary condition-related hypotheses using LMMs while accounting for repeated observations within participants.

### Methods S2

#### fNIRS data acquisition and analysis

##### Data acquisition

Cortical hemodynamic signals during face processing were recorded using a portable continuous-wave fNIRS system (NIRSport2, NIRx Medical Technologies, USA) at a sampling rate of 6.26 Hz. The system was equipped with 12 dual-wavelength emitters (760 nm, 850 nm) and 12 detectors, forming 28 measurement channels with a source-detector spacing of approximately 30 mm. Optodes were mounted on an elastic nylon cap according to the international 10–20 coordinate system to ensure stable scalp contact and consistent probe placement across participants (Table S1). According to the research purpose, the channels bilaterally covered the inferior frontal gyrus (IFG) and temporoparietal junction (TPJ) (see Figure S2C).

##### Data Preprocessing

fNIRS data were analyzed in MATLAB R2022b using the HOMER2 package (Huppert et al. 2009) and NIRS-SPM v4.1 (Ye et al. 2009). The complete analytical workflow is illustrated in Figure S2. First, poor-quality channels were automatically removed based on predefined criteria: raw intensity out of 10^-^³∼10⁵, SNR < 2.0, source– detector spacing exceeding 0–45 mm, or negative intensity values (Yucel et al. 2021). Valid signals were then converted to optical density (OD). Motion artifacts including transient spikes and baseline drifts were corrected via the Temporal Derivative Distribution Repair (TDDR) algorithm (Fishburn et al. 2019). This threshold-free automated method avoids subjective manual data rejection and retains data integrity. Corrected OD signals were converted to oxyhemoglobin (ΔHbO) and deoxyhemoglobin (ΔHbR) concentration changes using the modified Beer–Lambert law (MBLL). An age-matched differential pathlength factor (DPF = 2.5) was adopted for both wavelengths (Whiteman et al. 2018), tailored to our pediatric sample (mean age = 5.57 years).

##### Task-Related Brain Activity Quantification

Given the higher sensitivity of ΔHbO signals to regional cerebral blood flow and their stronger correlation with fMRI BOLD signals compared with ΔHbR, all task-evoked cortical activation analyses in this study were based on ΔHbO concentration changes.

A general linear model (GLM) was constructed to quantify event-related hemodynamic responses. Prior to model fitting, multi-dimensional noise reduction was performed: the wavelet minimum description length (Wavelet-MDL) algorithm was used to eliminate global signal baseline drift (Ye et al. 2009), and a pre-coloring filter based on the canonical hemodynamic response function (HRF) was applied to correct temporal autocorrelation and prevent statistical bias (Worsley and Friston 1995).

The GLM design matrix included four experimental task conditions and a unified baseline. Each 60-second task phase was modeled by convolving stimulus onset and duration with the canonical HRF. The baseline period covered all remaining recording time, including two 60-second resting blocks and all 10-second fixation trials. HRF temporal and dispersion derivatives were further incorporated into the model to accommodate individual and regional differences in hemodynamic response latency and peak amplitude.

After GLM parameter estimation, the canonical HRF beta weights (β-values) for each task condition were extracted. For subsequent statistical analysis, β-values were spatially averaged within the a priori defined regions of interest (ROIs; bilateral IFG and TPJ). Individual channel-level estimates within these ROIs were also retained to support supplementary fine-grained spatial analyses.

### Methods S3

#### Machine learning analyses approach

##### The Elastic Net regression model and settings

A data-driven, cross-validated regularized regression framework was adopted to predict children’s behavioral phenotypes based on task-evoked regional brain activation. To minimize information leakage, all preprocessing and model training procedures were performed exclusively within each training fold of the cross-validation procedure. Predictive features were derived from baseline-adjusted self-other activation contrasts of the predefined ROIs. Prior to model training, the SCQ scores were log-transformed, utilizing ln(*y* + 1), to mitigate potential skewness in the clinical distribution. Conversely, the ADOS comparison scores were not transformed and were retained in their original scale (y). To ensure computational reproducibility, the random number generator seed was strictly initialized for all partitioning and permutation steps.

An Elastic Net regression model with the mixing parameter fixed at α = 0.1 was used to integrate L1 (Lasso) variable selection and L2 (Ridge) regularization, enabling simultaneous feature selection and coefficient shrinkage. A 20-repeated five-fold cross-validation strategy was applied to improve model generalizability and reduce sampling variability. Importantly, all preprocessing procedures were conducted independently within each cross-validation training fold, with all parameters estimated solely from the training data.

Specifically, missing brain features and age values were imputed using training-set median values. Chronological age effects were then regressed out to isolate intrinsic predictive effects of neuroimaging features: age-related variance was estimated via linear models on the training data, and the resulting regression coefficients were subsequently applied to remove age effects from both the training and corresponding test data. All imaging features were standardized to zero mean and unit variance using statistics estimated from the training data, and the same transformation was subsequently applied to the corresponding test data.

Within each training fold, the regularization parameter (λ) was optimized using an internal three-fold cross-validation by minimizing the mean squared error (MSE), while the Elastic Net mixing parameter was fixed at α = 0.1 throughout the analysis. The optimized model was then applied to the held-out test fold to generate behavioral predictions. Five-fold cross-validation was repeated 20 times using different random data partitions. For each participant, predicted scores obtained across the repeated cross-validation runs were averaged to generate a single participant-level predicted score for subsequent model evaluation.

##### Model Evaluation and Feature Importance

Model predictive performance was evaluated using the R² indicator: Pearson’s or Spearman’s correlation coefficients between observed and predicted scores, cross-validated predictive coefficient of determination (predictive R²).

Feature importance was defined as the mean absolute regression weight across all cross-validation folds, and model stability was assessed via feature selection frequency. Features were ranked by importance to identify the core neural biomarkers corresponding to behavioral phenotypes.

##### Permutation Testing

A total of 1000 non-parametric permutation tests were performed to evaluate whether prediction performance exceeded chance level. In each permutation iteration, behavioral scores were randomly shuffled, and the complete machine learning pipeline was repeated identically to construct a null distribution of correlation coefficients. The empirical p-value was calculated separately for prediction correlation, predictive *R*^2^, and MAE as the proportion of permuted statistics that were at least as extreme as the observed statistic. To avoid zero p-values and effectively control Type I statistical error, p-values were calculated as (*N* + 1)/(1000 + 1), where *N* denotes the number of permutations exceeding the observed value (Phipson and Smyth 2010).

**Table S1.** Estimated MNI coordinates and corresponding anatomical regions for fNIRS channels.

| **Channel** | **MNI x** | **MNI y** | **MNI z** |
|---|---|---|---|
| Left inferior frontal gyrus |  |  |  |
| 1 | -37 | 63 | 3 |
| 2 | -50 | 46 | -11 |
| 3 | -42 | 56 | 15 |
| 4 | -55 | 38 | 0 |
| 6 | -58 | 26 | 16 |
| Left posterior superior temporal gyrus |  |  |  |
| 9 | -70 | -20 | 5 |
| Left posterior middle temporal gyrus |  |  |  |
| 10 | -71 | -34 | -11 |
| 12 | -69 | -48 | 8 |
| 13 | -64 | -60 | -6 |
| Left supramarginal gyrus |  |  |  |
| 11 | -69 | -33 | 27 |
| Left angular gyrus |  |  |  |
| 14 | -61 | -62 | 27 |
| Right inferior frontal gyrus |  |  |  |
| 15 | 41 | 63 | 4 |
| 16 | 53 | 45 | -11 |
| 17 | 45 | 55 | 16 |
| 18 | 57 | 38 | 1 |
| 20 | 61 | 25 | 17 |
| Right posterior superior temporal gyrus |  |  |  |
| 23 | 72 | -20 | 8 |
| Right posterior middle temporal gyrus |  |  |  |
| 24 | 72 | -36 | -9 |
| 26 | 69 | -50 | 11 |
| 27 | 64 | -61 | -5 |
| Right supramarginal gyrus |  |  |  |
| 25 | 70 | -34 | 29 |
| Right angular gyrus |  |  |  |
| 28 | 62 | -62 | 28 |

*Note.* MNI coordinates are probabilistic estimates generated via virtual spatial registration using NIRS-SPM, based on the standard 10-20 system geometry. The bilateral TPJ has been described as a functionally and anatomically heterogeneous region involving the posterior temporal cortex, including regions commonly referred to as posterior superior temporal gyrus (pSTG) and posterior middle temporal gyrus (pMTG), as well as inferior parietal regions, including angular gyrus (AG) and supramarginal gyrus (SMG).

**Table S2.** Linear mixed-effects models predicting hemodynamic responses across regions of interest in the TD group.

|  | Estimate (*SE*) | *t* | DF | *p* | 95%CI | Cohen’s *d* | Estimate (SE) | *t* | DF | *p* | 95%CI | Cohen’s *d* |
|---|---|---|---|---|---|---|---|---|---|---|---|---|
| **Fixed Effects** | **Left IFG** |  |  |  |  |  | **Right IFG** |  |  |  |  |  |
| (Intercept) | 1.322(0.245) | 5.404 | 25.847 | < .001 | [0.819, 1.825] | 1.063 | 0.857(0.186) | 4.608 | 112.000 | < .001 | [0.488, 1.226] | 0.435 |
| Face Type (Self = 1) | -0.312(0.175) | -1.785 | 82.631 | 0.078 | [-0.660, 0.036] | -0.196 | -0.244(0.186) | -1.312 | 112.000 | 0.192 | [-0.612, 0.125] | -0.124 |
| Red Dot (with RedDot=1) | -0.256(0.175) | -1.466 | 82.639 | 0.147 | [-0.604, 0.092] | -0.161 | 0.142(0.186) | 0.765 | 112.000 | 0.446 | [-0.226, 0.511] | 0.073 |
| Age | 0.234(0.188) | 1.248 | 25.606 | 0.223 | [-0.152, 0.620] | 0.247 | 0.094(0.143) | 0.653 | 112.000 | 0.515 | [-0.190, 0.378] | 0.062 |
| Face Type × Red Dot | 0.170(0.175) | 0.973 | 82.606 | 0.333 | [-0.178, 0.518] | 0.107 | 0.034(0.186) | 0.183 | 112.000 | 0.855 | [-0.334, 0.403] | 0.017 |
| **Random Effects** |  |  |  |  |  |  |  |  |  |  |  |  |
| Subject Variance | 0.878 |  |  |  |  |  | 0.000 |  |  |  |  |  |
| Residual Variance | 3.568 |  |  |  |  |  | 4.046 |  |  |  |  |  |
| **Model Fit** |  |  |  |  |  |  |  |  |  |  |  |  |
| Marginal R-squared | 0.061 |  |  |  |  |  | 0.024 |  |  |  |  |  |
| Conditional R-squared | 0.246 |  |  |  |  |  | 0.024 |  |  |  |  |  |
| **Fixed Effects** | **Left TPJ** |  |  |  |  |  | **Right TPJ** |  |  |  |  |  |
| (Intercept) | 0.959(0.294) | 3.256 | 28.395 | 0.003 | [0.356, 1.561] | 0.611 | 0.626(0.296) | 2.117 | 22.472 | 0.046 | [0.013, 1.239] | 0.447 |
| Face Type (Self = 1) | 0.040(0.210) | 0.191 | 85.914 | 0.849 | [-0.378, 0.458] | 0.021 | -0.516(0.219) | -2.352 | 78.255 | 0.021 | [-0.952, -0.079] | -0.266 |
| Red Dot (with RedDot=1) | 0.124(0.210) | 0.589 | 85.915 | 0.558 | [-0.294, 0.542] | 0.064 | -0.039(0.219) | -0.180 | 78.255 | 0.858 | [-0.476, 0.397] | -0.020 |
| Age | 0.569(0.227) | 2.503 | 28.813 | 0.018 | [0.104, 1.034] | 0.466 | 0.336(0.233) | 1.443 | 23.439 | 0.162 | [-0.145, 0.816] | 0.298 |
| Face Type × Red Dot | -0.060(0.210) | -0.283 | 85.914 | 0.778 | [-0.478, 0.359] | -0.030 | 0.542(0.219) | 2.470 | 78.255 | 0.016 | [0.105, 0.978] | 0.279 |
| **Random Effects** |  |  |  |  |  |  |  |  |  |  |  |  |
| Subject Variance | 1.272 |  |  |  |  |  | 1.160 |  |  |  |  |  |
| Residual Variance | 5.207 |  |  |  |  |  | 5.564 |  |  |  |  |  |
| **Model Fit** |  |  |  |  |  |  |  |  |  |  |  |  |
| Marginal R-squared | 0.080 |  |  |  |  |  | 0.100 |  |  |  |  |  |
| Conditional R-squared | 0.260 |  |  |  |  |  | 0.255 |  |  |  |  |  |
| **Simple Effects for Interaction** |  |  |  |  |  |  |  |  |  |  |  |  |
| Simple Self (at NoRed) |  |  |  |  |  |  |  | -0.084 |  | 0.934 |  |  |
| Simple Self (at RedDot) |  |  |  |  |  |  |  | 3.406 |  | 0.001 |  |  |

**Table S3.**
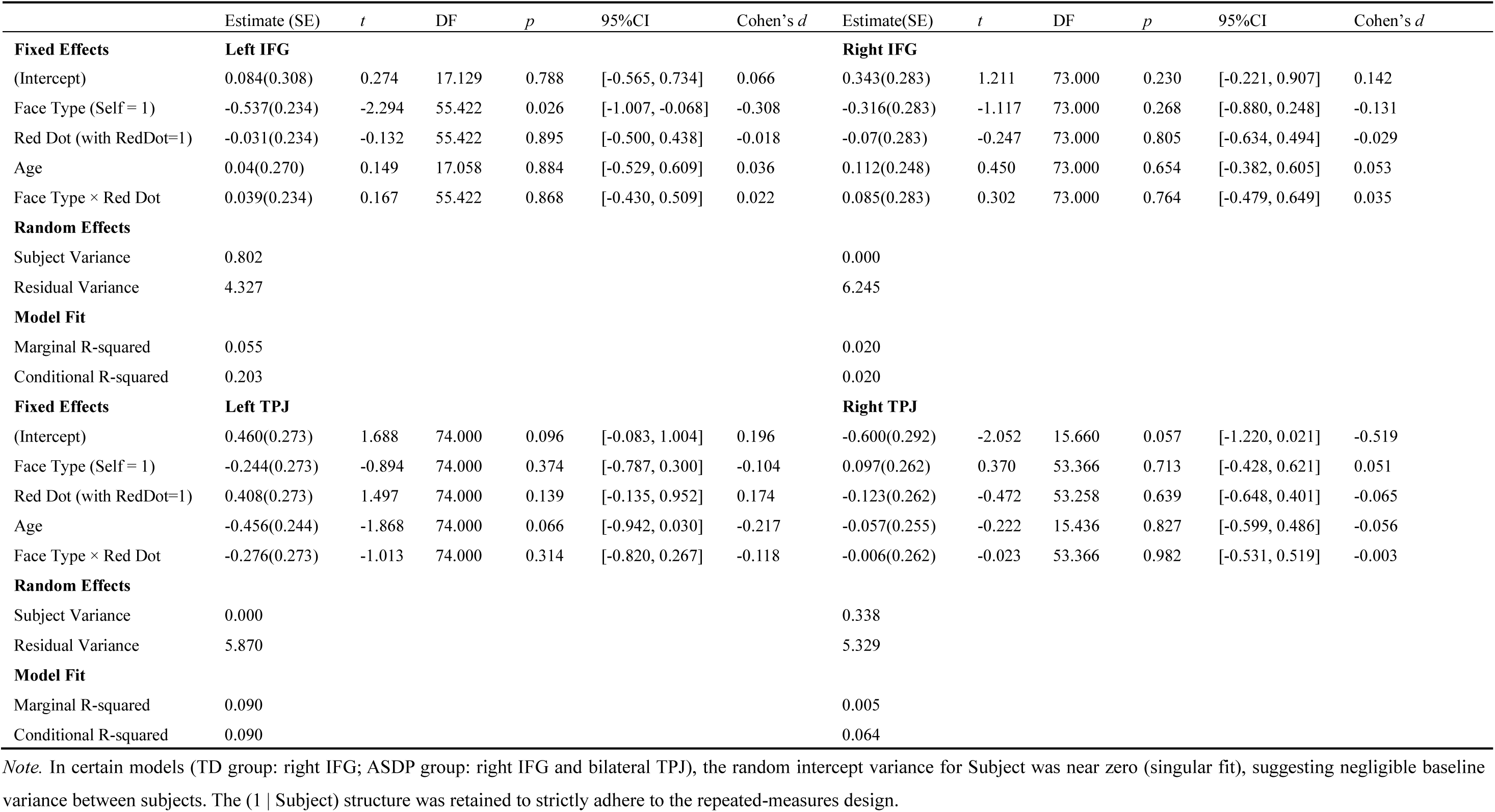
Linear mixed-effects models predicting hemodynamic responses across regions of interest in the ASDP group.

**Table S4.** Linear mixed-effects models predicting hemodynamic responses across regions of interest in the ASDF group.

|  | Estimate (SE) | *t* | DF | *p* | 95%CI | Cohen’s *d* | Estimate(SE) | *t* | DF | *p* | 95%CI | Cohen’s *d* |
|---|---|---|---|---|---|---|---|---|---|---|---|---|
| **Fixed Effects** | **Left IFG** |  |  |  |  |  | **Right IFG** |  |  |  |  |  |
| (Intercept) | 0.196(0.230) | 0.852 | 28.268 | 0.401 | [-0.275, 0.666] | 0.160 | -0.016(0.190) | -0.084 | 26.670 | 0.934 | [-0.406, 0.374] | -0.016 |
| Face Type (Self = 1) | -0.068(0.157) | -0.433 | 85.753 | 0.666 | [-0.380, 0.244] | -0.047 | -0.261(0.167) | -1.558 | 85.478 | 0.123 | [-0.594, 0.072] | -0.168 |
| Red Dot (with RedDot=1) | -0.124(0.157) | -0.788 | 85.800 | 0.433 | [-0.436, 0.189] | -0.085 | 0.167(0.167) | 0.996 | 84.015 | 0.322 | [-0.166, 0.499] | 0.109 |
| Age | 0.176(0.158) | 1.115 | 28.161 | 0.274 | [-0.147, 0.499] | 0.210 | -0.035(0.130) | -0.268 | 26.652 | 0.791 | [-0.303, 0.233] | -0.052 |
| Face Type × Red Dot | 0.084(0.157) | 0.537 | 85.800 | 0.593 | [-0.228, 0.397] | 0.058 | 0.124(0.167) | 0.741 | 84.015 | 0.461 | [-0.209, 0.457] | 0.081 |
| **Random Effects** |  |  |  |  |  |  |  |  |  |  |  |  |
| Subject Variance | 0.845 |  |  |  |  |  | 0.241 |  |  |  |  |  |
| Residual Variance | 2.903 |  |  |  |  |  | 3.300 |  |  |  |  |  |
| **Model Fit** |  |  |  |  |  |  |  |  |  |  |  |  |
| Marginal R-squared | 0.025 |  |  |  |  |  | 0.031 |  |  |  |  |  |
| Conditional R-squared | 0.244 |  |  |  |  |  | 0.097 |  |  |  |  |  |
| **Fixed Effects** | **Left TPJ** |  |  |  |  |  | **Right TPJ** |  |  |  |  |  |
| (Intercept) | 0.004(0.243) | 0.017 | 28.428 | 0.986 | [-0.492, 0.501] | 0.003 | -0.079(0.195) | -0.407 | 109.000 | 0.685 | [-0.466, 0.307] | -0.039 |
| Face Type (Self = 1) | -0.220(0.186) | -1.184 | 86.021 | 0.240 | [-0.589, 0.149] | -0.128 | -0.301(0.195) | -1.544 | 109.000 | 0.125 | [-0.688, 0.085] | -0.148 |
| Red Dot (with RedDot=1) | -0.188(0.186) | -1.013 | 86.027 | 0.314 | [-0.557, 0.181] | -0.109 | -0.173(0.195) | -0.884 | 109.000 | 0.379 | [-0.559, 0.214] | -0.085 |
| Age | -0.124(0.167) | -0.743 | 28.740 | 0.463 | [-0.466, 0.218] | -0.139 | -0.038(0.131) | -0.292 | 109.000 | 0.771 | [-0.298, 0.221] | -0.028 |
| Face Type × Red Dot | 0.149(0.186) | 0.802 | 86.027 | 0.425 | [-0.22, 0.518] | 0.086 | 0.016(0.195) | 0.080 | 109.000 | 0.936 | [-0.371, 0.402] | 0.008 |
| **Random Effects** |  |  |  |  |  |  |  |  |  |  |  |  |
| Subject Variance | 0.731 |  |  |  |  |  | 0.000 |  |  |  |  |  |
| Residual Variance | 4.057 |  |  |  |  |  | 4.340 |  |  |  |  |  |
| **Model Fit** |  |  |  |  |  |  |  |  |  |  |  |  |
| Marginal R-squared | 0.029 |  |  |  |  |  | 0.028 |  |  |  |  |  |
| Conditional R-squared | 0.177 |  |  |  |  |  | 0.028 |  |  |  |  |  |

**Table S5.** The scores of Social Communication Questionnaire across three groups.

|  | **TD (N=30)** | **ASD (N=50)** | **ASDP (N=20)** | **ASDF (N=30)** | **TD vs ASD** | **ASDP vs ASDF** |
|---|---|---|---|---|---|---|
| Total Score | 5.73 ± 3.18 (n=30) | 20.55 ± 6.21 (n=49) | 21.37 ± 6.79 (n=19) | 20.03 ± 5.87 (n=30) | *** | *** |
| Reciprocal Social Interaction | 0.70 ± 0.92 (n=30) | 9.08 ± 3.01 (n=49) | 9.42 ± 3.56 (n=19) | 8.87 ± 2.65 (n=30) | *** | *** |
| Communication | 2.00 ± 1.20 (n=30) | 6.73 ± 2.37 (n=49) | 6.58 ± 2.67 (n=19) | 6.83 ± 2.20 (n=30) | *** | *** |
| Restricted, Repetitive, and Stereotyped Behaviors | 2.77 ± 1.81 (n=30) | 4.31 ± 2.35 (n=49) | 4.84 ± 2.34 (n=19) | 3.97 ± 2.33 (n=30) | ** | ** |

*Note.* Asterisks indicate uncorrected p values: \*p < 0.05, \*\*p < 0.01, and \*\*\*p < 0.001

## Supplemental figures

**Figure S1:**
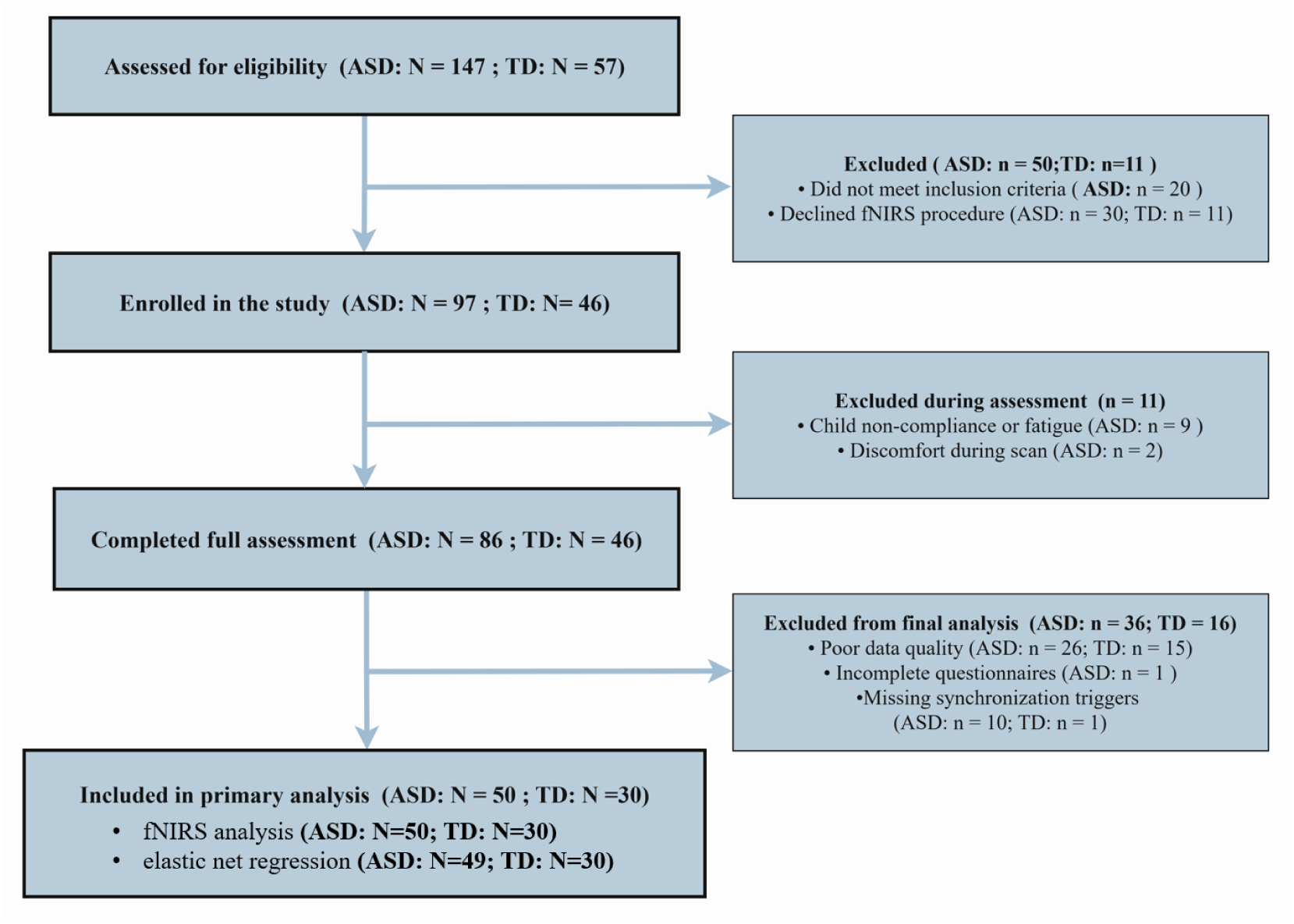
Participant flow diagram. A total of 147 ASD and 57 TD children were assessed; after exclusions (e.g., ineligibility, refusal, poor compliance, data quality issues), 50 ASD and 30 TD were included in the analysis.

**Figure S2:**
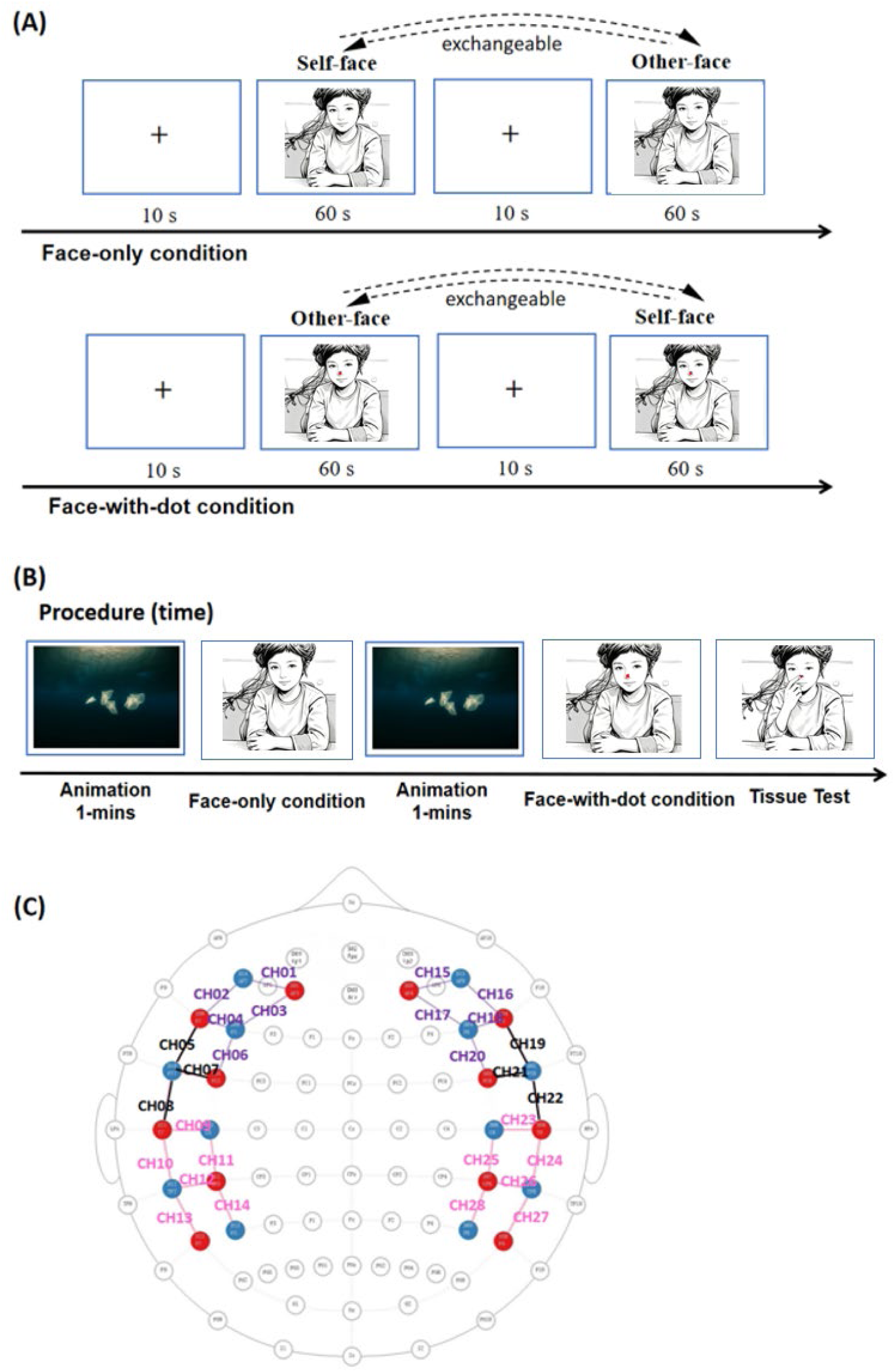
Experimental task and procedure. (A) Participants performed two task conditions in a face-processing paradigm: a face-only condition and a face-with-red-dot condition. (B) Following the experimental setup, participants completed the face-without-red-dot task, the face-with-red-dot task, and an additional tissue test. (C) fNIRS channels were placed over the bilateral inferior frontal gyrus (IFG) (purple lines) and temporoparietal junction (TPJ) (pink lines).

**Figure S3:**
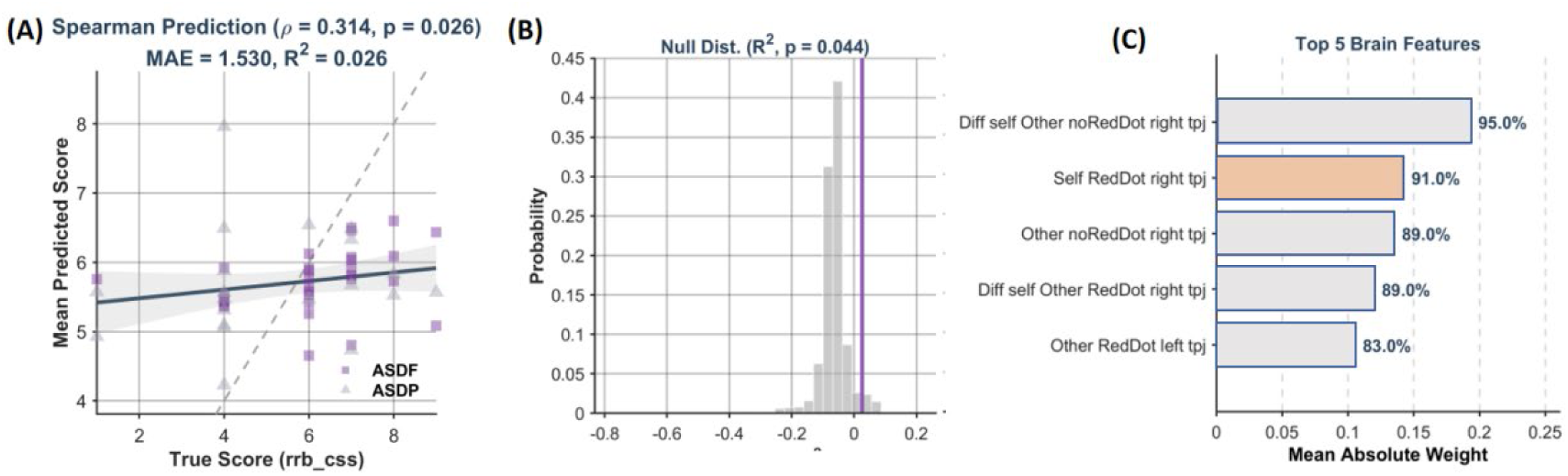
Elastic Net Regression Analysis of Brain Activation Predicting RRB CSS scores in ASD. Neural activity during face processing significantly predicted ADOS scores in individuals with ASD (A). This predictive effect survived permutation testing (B). The top five contributing brain features were further extracted and visualized in a panel that included the bodily-self-related right TPJ (yellow bar).

